# *cytoviewer:* an R/Bioconductor package for interactive visualization and exploration of highly multiplexed imaging data

**DOI:** 10.1101/2023.05.24.542115

**Authors:** Lasse Meyer, Nils Eling, Bernd Bodenmiller

**Affiliations:** Department of Quantitative Biomedicine, University of Zurich, Zurich, Switzerland; Institute for Molecular Health Sciences, ETH Zurich, Zurich, Switzerland; Life Science Zurich Graduate School, ETH Zurich and University of Zurich, Zurich, Switzerland

## Abstract

**Summary:** Highly multiplexed imaging enables single-cell-resolved detection of numerous biological molecules in their spatial tissue context. Interactive data visualization of multiplexed imaging data is necessary for quality control and hypothesis examination. Here, we describe *cytoviewer*, an R/Bioconductor package for interactive visualization and exploration of multi-channel images and segmentation masks. The *cytoviewer* package supports flexible generation of image composites, allows side-by-side visualization of single channels, and facilitates the spatial visualization of single-cell data in the form of segmentation masks. The package operates on *SingleCellExperiment, SpatialExperiment* and *CytoImageList* objects and therefore integrates with the Bioconductor framework for single-cell and image analysis. Users of *cytoviewer* need little coding expertise, and the graphical user interface allows user-friendly navigation. We showcase the functionality of *cytoviewer* by analysis of an imaging mass cytometry dataset of cancer patients.

**Availability:** The *cytoviewer* package can be installed from Bioconductor via https://www.bioconductor.org/packages/release/bioc/html/cytoviewer.html. The development version and further instructions can be found on GitHub at https://github.com/BodenmillerGroup/cytoviewer. We provide an R script to exemplify the usage of *cytoviewer* in the supplementary information.

**Supplementary informations:** Supplementary data are available online.

## Introduction

Highly multiplexed imaging allows spatially and single-cell-resolved detection of dozens of biological molecules, including proteins and nucleic acids, *in situ*. These technologies facilitate an in-depth analysis of complex systems and diseases such as the tumor microenvironment (Hoch et al., 2022; Jackson et al., 2020; Risom et al., 2022) and type 1 diabetes progression (Damond et al., 2019). Imaging-based spatial proteomics methods (Moffitt et al., 2022) can be broadly divided into fluorescent cyclic approaches such as tissue-based cyclic immunofluorescence (t-CyCIF) (Lin et al., 2018) and one-step masstag based approaches that include multiplexed ion beam imaging (MIBI) (Angelo et al., 2014) and imaging mass cytometry (IMC) (Giesen et al., 2014).

To fully leverage the information contained in multiplexed imaging data, computational tools are necessary. The main analysis steps, irrespective of the biological question, are 1) image quality control, 2) image pre-processing and segmentation, and 3) single-cell and spatial analysis (Windhager et al., 2021). Interactive image visualization greatly benefits image and segmentation quality control and hypothesis generation and verification. However, commonly used programs, such as histoCAT (Schapiro et al., 2017), QuPath (Bankhead et al., 2017), and others (Schindelin et al., 2012; Somarakis et al., 2021), have little interoperability with other frameworks and programming languages. The recently developed napari image viewer, which operates in Python, bridges the gap between multiplexed image visualization and data analysis (Chiu et al., 2022), but similar tools that operate in the statistical programming language R have not been developed.

Here, we present the R/Bioconductor *cytoviewer* package for interactive multi-channel image and segmentation mask visualization in R. The *cytoviewer* package builds on the *cytomapper* R/Bioconductor package (Eling et al., 2020) and extends its static visualization abilities via an interactive and user-friendly *shiny* application It provides interactive visualization strategies in a similar fashion as the *iSEE* package (Rue-Albrecht et al., 2018) offers for single-cell data and can be seamlessly harmonized with any step of the data analysis workflow in R. Users can overlay individual images with segmentation masks, visualize cell-specific metadata, and download generated images. The *cytoviewer* package integrates into the Bioconductor framework (Gentleman et al., 2004) for single-cell and image analysis leveraging the image handling and analysis strategies from the *EBImage* Bioconductor package (Pau et al., 2010) and building on commonly used Bioconductor classes such as *SingleCellExperiment, SpatialExperiment* (Amezquita et al., 2020; Righelli et al., 2022), and *CytoImageList* (Eling et al., 2020). We showcase the functionality and potential application fields of *cytoviewer* by demonstrating visual exploration of an IMC dataset of cancer patients.

## Results

The R/Bioconductor *cytoviewer* package leverages the reactive programming framework of the popular R *shiny* and *shinydashboard* packages (Jia et al., 2022), is cross-platform compatible, and launches an interactive web application. The graphical user interface of the *cytoviewer* package has three main parts: body, sidebar, and header **(Figure 1A)**. The body of *cytoviewer* features the image viewer. The viewer can switch between image-level visualization, which shows the pixel-level intensities of all selected markers either combined (Composite) or separately (Channels), and cell-level visualization, which displays cell-level information on segmentation masks (Masks) **(Supplementary Note S1.1)**. Controls for sample selection and image and mask visualization settings as well as image appearance/filters are found in the sidebar menu. The header section contains the package version, R session information, a help page, and a drop-down menu for image downloads.

**Figure 1:**
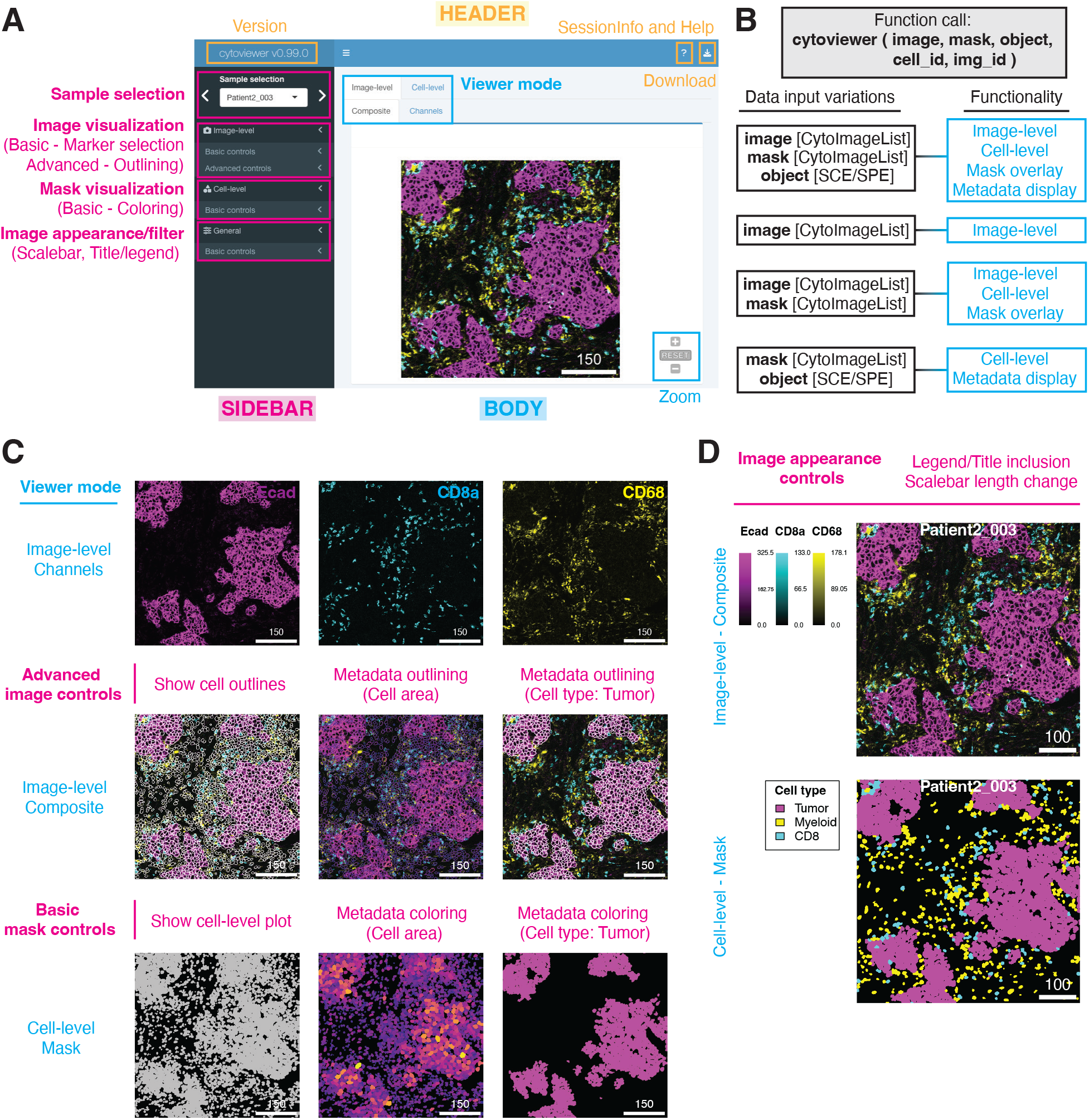
*cytoviewer* interface and functionality. **(A)** The graphical user interface of *cytoviewer* is divided into a body, header, and sidebar. The body of *cytoviewer* includes the image viewer, which has three tabs: Composite (Image-level), Channels (Image-level), and Mask (Cell-level). Zooming is supported for Composite and Mask tabs. The package version, R session information, help page, and a drop-down menu for image downloads are located in the header. The sidebar menu has controls for sample selection, image visualization, mask visualization, and general settings. In the image shown, the scale bar is 150 *µ*m. **(B)** The supported functionality (right) of *cytoviewer* depends on the data inputs provided (left). To match information between the objects, cell (cell_id) and image (img_id) identifiers can be provided. SCE/SPE = SingleCellExperiment/SpatialExperiment. **(C)** *cytoviewer* supports different viewing modes. Top: The “channels” tab of image-level visualization displays selected channels in individual images. Shown are Ecad (magenta), CD8a (cyan), and CD68 (yellow) marking tumor cells, CD8^+^ T cells, and myeloid cells, respectively. Center: The “composite” tab of image-level visualization can be overlayed with cell outlines, which can be colored by cell-specific metadata. Shown here are cell area (continuous; plasma) and cell type (categorical; tumor cells in white) information. Channel color settings are as follows for all markers: Contrast: 2,5; Brightness: 1; Gamma: 1.2. Bottom: The “mask” tab can be used to visualize segmentation mask outlines that can be colored by cell-specific metadata. Shown here are cell area (continuous; plasma) and cell type (categorical; tumor cells in magenta) information. Scale bars in all images are 150 *µ*m. **(D)** “Image appearance” controls can be used to add legends or titles and to change the scale bar length for image-level (top) and cell level (bottom) visualization. The cell-level mask plot shown depicts tumor (magenta), myeloid (yellow), and CD8^+^ T cells (cyan). Scale bars are 100 *µ*m.

The *cytoviewer* function call takes up to five arguments **(Figure 1B)**. Images must be provided as a *CytoImageList* object containing one or multiple multi-channel images where each channel represents the pixel intensities of one marker. Segmentation masks in *CytoImageList* format can be added if desired. Segmentation masks are represented as single-channel images containing integer values for cells or zero for background. Furthermore, *SingleCellExperiment* or *SpatialExperiment* class objects can be provided to allow single-cell specific metadata visualization. The full functionality of *cytoviewer* is leveraged when images, segmentation masks, and a metadata object are provided **(Figure 1B)**. This allows comprehensive image-level and cell-level visualization, enables image overlays with segmentation masks, and cell-specific metadata visualization.

To demonstrate the functionality and potential applications of *cytoviewer*, we explored an example IMC dataset from the Integrated iMMUnoprofiling of large adaptive CANcer patient cohort project (immucan.eu) **(Supplementary Note S1.2)**. For IMC, tissue sections are stained with antibodies tagged with isotopically pure rare earth metals, the tissue is laser ablated, and tags are detected by mass spectrometry to produce high-dimensional images (Giesen et al., 2014). Here, we demonstrate the different viewing modes of *cytoviewer* by analyses of images from a breast cancer patient (Patient2_003) **(Figure 1C, Supplementary Figure S1)**.

Image visualization control is split into basic and advanced control modes. Basic controls support the selection of up to six channels with separate color control settings for each (contrast, brightness, gamma, and channel color). In the example shown here, we visualized expression of *Ecad, CD8a*, and *CD68*, which are markers for epithelial and tumor cells, CD8^+^ T cells, and myeloid cells, respectively **(Figure 1A, Figure 1C -*top*)**. This image visualization step can support qualitative assessment of signal sensitivity and specificity.

In the advanced image control mode, the user can choose to overlay the displayed images with provided segmentation masks **(Figure 1C – *center*)**. Outline color and thickness can be adjusted by the user. Of note, this step can support evaluation of cell segmentation quality, which is essential for downstream data analysis. Moreover, the masks can be outlined by cell-specific metadata from the *SingleCellExperiment/SpatialExperiment* object. For categorical and continuous metadata entries, the user can choose between discrete colors and continuous color palettes (viridis, inferno, plasma), respectively. By outlining the masks with the cell area and cell type information (e.g., tumor), correct phenotype assignment can be visually confirmed (e.g., tumor cells are Ecad^+^ and tumor cells have larger areas than other cells).

The user can decide to display the provided segmentation masks **(Figure 1C – *bottom*)**. Coloring of the masks by cell-specific metadata (categorical and continuous) is possible and can be used for visual assessment of, for example, tumor cell areas and structures.

Using image appearance controls, the user can adjust the scale bar length and include legends or image titles. These features can be used for image-level and cell-level visualization and can aid in interpretation of phenotype co-localization such as infiltration of CD8^+^ T cells into the tumor core **(Figure 1D)**. Furthermore, from the image filters section, the user can control pixel-wise interpolation (default) and apply Gaussian filters on the image-level **(Supplementary Figure S2)**.

The *cytoviewer* package also supports rapid download of the generated images in publication quality. For download, the user specifies a file name, selects the image of interest (Composite, Channels, Mask) and the file format (pdf, png) **(Supplementary Note S1.3)**.

## Supporting information

R script

## Conclusion

The *cytoviewer* package provides a versatile and easy-to-use graphical user interface for interactive visualization of highly multiplexed imaging data in R. *cytoviewer* is accessible to researchers with little bioinformatics training and can support every step of the highly multiplexed data analysis workflow in R (Windhager et al., 2021) including visual cell-segmentation quality control, cell phenotype confirmation, and hypothesis examination. Here, we demonstrated the use of *cytoviewer* by exploring IMC data. However, data from other technologies such as t-CyCIF (Lin et al., 2018) or MIBI (Angelo et al., 2014), which produce pixel-level intensities and (optionally) segmentation masks, can be interactively visualized with *cytoviewer* as long as the input format is appropriate. The *cytoviewer* package, together with the related *cytomapper* package (Eling et al., 2020), are a well-integrated R/Bioconductor toolbox for highly multiplexed imaging data visualization in R relying on data containers such as *SingleCellExperiment/SpatialExperiment* (Amezquita et al., 2020; Righelli et al., 2022) and *CytoImageList* (Eling et al., 2020).

We envision that the *cytoviewer* package will meet the needs of the fast-growing community of highly multiplexed imaging users (Hickey et al., 2022) by providing user-friendly and rich data visualization that seamlessly integrates and supports the highly multiplexed imaging data analysis workflow in R.

## References

Amezquita, R. A., Lun, A. T. L., Becht, E., Carey, V. J., Carpp, L. N., Geistlinger, L., Marini, F., Rue-Albrecht, K., Risso, D., Soneson, C., Waldron, L., Pagès, H., Smith, M. L., Huber, W., Morgan, M., Gottardo, R., & Hicks, S. C. (2020). Orchestrating single-cell analysis with Bioconductor. Nature Methods, 17(2), 137–145. https://doi.org/10.1038/s41592-019-0654-x

Angelo, M., Bendall, S. C., Finck, R., Hale, M. B., Hitzman, C., Borowsky, A. D., Levenson, R. M., Lowe, J. B., Liu, S. D., Zhao, S., Natkunam, Y., & Nolan, G. P. (2014). Multiplexed ion beam imaging of human breast tumors. Nature Medicine, 20(4), 436–442. https://doi.org/10.1038/nm.3488

Bankhead, P., Loughrey, M. B., Fernández, J. A., Dombrowski, Y., McArt, D. G., Dunne, P. D., McQuaid, S., Gray, R. T., Murray, L. J., Coleman, H. G., James, J. A., Salto-Tellez, M., & Hamilton, P. W. (2017). QuPath: Open source software for digital pathology image analysis. Scientific Reports, 7(1), 16878. https://doi.org/10.1038/s41598-017-17204-5

Chiu, C.-L., Clack, N., & community, the napari. (2022). napari: a Python Multi-Dimensional Image Viewer Platform for the Research Community. Microscopy and Microanalysis, 28(S1), 1576–1577. https://doi.org/10.1017/S1431927622006328

Damond, N., Engler, S., Zanotelli, V. R. T., Schapiro, D., Wasserfall, C. H., Kusmartseva, I., Nick, H. S., Thorel, F., Herrera, P. L., Atkinson, M. A., & Bodenmiller, B. (2019). A Map of Human Type 1 Diabetes Progression by Imaging Mass Cytometry. Cell Metabolism, 29(3), 755-768.e5. https://doi.org/10.1016/j.cmet.2018.11.014

Eling, N., Damond, N., Hoch, T., & Bodenmiller, B. (2020). cytomapper: an R/Bioconductor package for visualization of highly multiplexed imaging data. Bioinformatics, 36(24), 5706–5708. https://doi.org/10.1093/bioinformatics/btaa1061

Gentleman, R. C., Carey, V. J., Bates, D. M., Bolstad, B., Dettling, M., Dudoit, S., Ellis, B., Gautier, L., Ge, Y., Gentry, J., Hornik, K., Hothorn, T., Huber, W., Iacus, S., Irizarry, R., Leisch, F., Li, C., Maechler, M., Rossini, A. J., … Zhang, J. (2004). Bioconductor: open software development for computational biology and bioinformatics. Genome Biology, 5(10), R80. https://doi.org/10.1186/gb-2004-5-10-r80

Giesen, C., Wang, H. A. O., Schapiro, D., Zivanovic, N., Jacobs, A., Hattendorf, B., Schüffler, P. J., Grolimund, D., Buhmann, J. M., Brandt, S., Varga, Z., Wild, P. J., Günther, D., & Bodenmiller, B. (2014). Highly multiplexed imaging of tumor tissues with subcellular resolution by mass cytometry. Nature Methods, 11(4), 417–422. https://doi.org/10.1038/nmeth.2869

Hickey, J. W., Neumann, E. K., Radtke, A. J., Camarillo, J. M., Beuschel, R. T., Albanese, A., McDonough, E., Hatler, J., Wiblin, A. E., Fisher, J., Croteau, J., Small, E. C., Sood, A., Caprioli, R. M., Angelo, R. M., Nolan, G. P., Chung, K., Hewitt, S. M., Germain, R. N., … Saka, S. K. (2022). Spatial mapping of protein composition and tissue organization: a primer for multiplexed antibody-based imaging. Nature Methods, 19(3), 284–295. https://doi.org/10.1038/s41592-021-01316-y

Hoch, T., Schulz, D., Eling, N., Gómez, J. M., Levesque, M. P., & Bodenmiller, B. (2022). Multiplexed imaging mass cytometry of the chemokine milieus in melanoma characterizes features of the response to immunotherapy. Science Immunology, 7(70), eabk1692. https://doi.org/10.1126/sciimmunol.abk1692

Jackson, H. W., Fischer, J. R., Zanotelli, V. R. T., Ali, H. R., Mechera, R., Soysal, S. D., Moch, H., Muenst, S., Varga, Z., Weber, W. P., & Bodenmiller, B. (2020). The single-cell pathology landscape of breast cancer. Nature, 578(7796), 615–620. https://doi.org/10.1038/s41586-019-1876-x

Jia, L., Yao, W., Jiang, Y., Li, Y., Wang, Z., Li, H., Huang, F., Li, J., Chen, T., & Zhang, H. (2022). Development of interactive biological web applications with R/Shiny. Briefings in Bioinformatics, 23(1), bbab415. https://doi.org/10.1093/bib/bbab415

Lin, J.-R., Izar, B., Wang, S., Yapp, C., Mei, S., Shah, P. M., Santagata, S., & Sorger, P. K. (2018). Highly multiplexed immunofluorescence imaging of human tissues and tumors using t-CyCIF and conventional optical microscopes. ELife, 7, e31657. https://doi.org/10.7554/eLife.31657

Moffitt, J. R., Lundberg, E., & Heyn, H. (2022). The emerging landscape of spatial profiling technologies. Nature Reviews Genetics, 23(12), 741–759. https://doi.org/10.1038/s41576-022-00515-3

Pau, G., Fuchs, F., Sklyar, O., Boutros, M., & Huber, W. (2010). EBImage—an R package for image processing with applications to cellular phenotypes. Bioinformatics, 26(7), 979–981. https://doi.org/10.1093/bioinformatics/btq046

Righelli, D., Weber, L. M., Crowell, H. L., Pardo, B., Collado-Torres, L., Ghazanfar, S., Lun, A. T. L., Hicks, S. C., & Risso, D. (2022). SpatialExperiment: infrastructure for spatially-resolved transcriptomics data in R using Bioconductor. Bioinformatics, 38(11), 3128–3131. https://doi.org/10.1093/bioinformatics/btac299

Risom, T., Glass, D. R., Averbukh, I., Liu, C. C., Baranski, A., Kagel, A., McCaffrey, E. F., Greenwald, N. F., Rivero-Gutiérrez, B., Strand, S. H., Varma, S., Kong, A., Keren, L., Srivastava, S., Zhu, C., Khair, Z., Veis, D. J., Deschryver, K., Vennam, S., … Angelo, M. (2022). Transition to invasive breast cancer is associated with progressive changes in the structure and composition of tumor stroma. Cell, 185(2), 299-310.e18. https://doi.org/10.1016/j.cell.2021.12.023

Rue-Albrecht, K., Marini, F., Soneson, C., & Lun, A. T. L. (2018). iSEE: Interactive SummarizedExperiment Explorer [version 1; peer review: 3 approved]. F1000Research, 7(741). https://doi.org/10.12688/f1000research.14966.1

Schapiro, D., Jackson, H. W., Raghuraman, S., Fischer, J. R., Zanotelli, V. R. T., Schulz, D., Giesen, C., Catena, R., Varga, Z., & Bodenmiller, B. (2017). histoCAT: analysis of cell phenotypes and interactions in multiplex image cytometry data. Nature Methods, 14(9), 873–876. https://doi.org/10.1038/nmeth.4391

Schindelin, J., Arganda-Carreras, I., Frise, E., Kaynig, V., Longair, M., Pietzsch, T., Preibisch, S., Rueden, C., Saalfeld, S., Schmid, B., Tinevez, J.-Y., White, D. J., Hartenstein, V., Eliceiri, K., Tomancak, P., & Cardona, A. (2012). Fiji: an open-source platform for biological-image analysis. Nature Methods, 9(7), 676–682. https://doi.org/10.1038/nmeth.2019

Somarakis, A., Unen, V. Van, Koning, F., Lelieveldt, B., & Höllt, T. (2021). ImaCytE: Visual Exploration of Cellular Micro-Environments for Imaging Mass Cytometry Data. IEEE Transactions on Visualization and Computer Graphics, 27(1), 98–110. https://doi.org/10.1109/TVCG.2019.2931299

Windhager, J., Bodenmiller, B., & Eling, N. (2021). An end-to-end workflow for multiplexed image processing and analysis. BioRxiv, 2021.11.12.468357. https://doi.org/10.1101/2021.11.12.468357

