## Supplementary Information for "*cytoviewer:* an R/Bioconductor package for interactive visualization and exploration of highly multiplexed imaging data"

Table of Content:

S1 Supplementary Notes:

S1.1 Function usage of different viewing modes

S1.2 The example IMC cancer dataset

S1.3 Download options

S2 Code and data availability

S3 Supplementary Figures:

Supplemental Figure S1 cytoviewer GUI overview

Supplemental Figure S2 cytoviewer filter overview

S4 References

### S1 Supplementary Notes

#### S1.1 Function usage and defaults for different viewing modes

The *cytoviewer* package builds on the R/Bioconductor *cytomapper* package (Eling et al., 2020) and utilizes its functions and data containers. Image-level visualization of the *cytoviewer* package is based on the *plotPixels* function of the R/Bioconductor *cytomapper* package (Eling et al., 2020). For more information, please refer to the help page found at *?cytomapper::plotPixels* and to the *cytomapper* package vignette.

In *cytoviewer*, basic controls for image-level visualization allow selection of up to six markers/channels with the following default colors: magenta, cyan, yellow, red, green, and blue. Colors are scaled between the minimum and maximum pixel intensities across the displayed image. Therefore, when selecting images of different samples, the range of pixel intensities can change. Advanced image controls support the overlay of images with segmentation masks (default color: white). When outlining the masks by cell-specific metadata, categorical and continuous metadata entries are colored using discrete colors (from the *RColorBrewer::brewer_pal()* function) and continuous color palettes (viridis, inferno, plasma from the *viridis* package). The "composite" tab of image-level visualization has zoom-controls.

Cell-level visualization of the *cytoviewer* package is based on the *plotCells* function of the R/Bioconductor *cytomapper* package (Eling et al., 2020). For more information, please refer to the help page found at *?cytomapper::plotCells* and to the *cytomapper* package vignette. Basic controls for cell-level visualization allow display (default color: gray) and coloring of masks by cell-specific metadata. Akin to image-level visualization, categorical and continuous metadata entries are colored using discrete colors (from the *RColorBrewer::brewer_pal()* function) and continuous color palettes (viridis, inferno, plasma from the *viridis* package). The "masks" tab of cell-level visualization has zoom-controls.

#### S1.2 The example IMC cancer dataset

The IMC dataset used for *cytoviewer* demonstration was generated as part of the Integrated iMMUnoprofiling of large adaptive CANcer patient cohort project (immucan.eu) with the Hyperion instrument from Standard BioTools (https://www.standardbio.com/products/instruments/hyperion). The dataset includes images of samples from four cancer patients diagnosed with different tumor types (head and neck cancer, breast cancer, lung cancer and colorectal cancer). For this demonstration, images from a breast cancer patient (Patient2_003) were used.

The data input objects were processed with the IMC data analysis workflow (https://bodenmillergroup.github.io/IMCDataAnalysis/) using functionality from the *steinbock* framework (Windhager et al., 2021) and the *imcRtools* package (https://github.com/BodenmillerGroup/imcRtools) among others. Data were downloaded from https://zenodo.org/record/7647079. The image data were stored as a *CytoImageList* object containing the spillover corrected multi-channel images, and the mask *CytoImageList* object was used to store single-channel segmentation masks. The single-cell data were stored in *SpatialExperiment* format (Righelli et al., 2022). Metadata information generated during the analysis was stored in the *colData* slot. For more details, please refer to https://bodenmillergroup.github.io/IMCDataAnalysis/.

#### S1.3 Download options

Image download controls are part of the header section as a drop-down menu. The user can specify a file name and select the image of interest (Composite, Channels, Mask) and the file format (pdf, png). When the download button is clicked, a pop-up window will appear where the user can specify the download location on the local machine. Individual images from the channels viewing mode are downloaded as a *.zip* file.

### S2 Code and data availability

All analysis were performed using Bioconductor 3.17, R version 4.3.0 and *cytoviewer* version 1.0.0.

All analysis code and instructions for data analysis are available online at:
*cytoviewer_publication_analysis.Rmd*

The *cytoviewer* package can be installed from Bioconductor:
https://www.bioconductor.org/packages/release/bioc/html/cytoviewer.html

The development version of *cytoviewer* can be found on GitHub:
https://github.com/BodenmillerGroup/cytoviewer

A static website with further instructions on package usage and functionality can be found at:
https://bodenmillergroup.github.io/cytoviewer/

The example IMC dataset used for the present publication is available at: https://zenodo.org/record/7647079

### S3 Supplementary Figures:

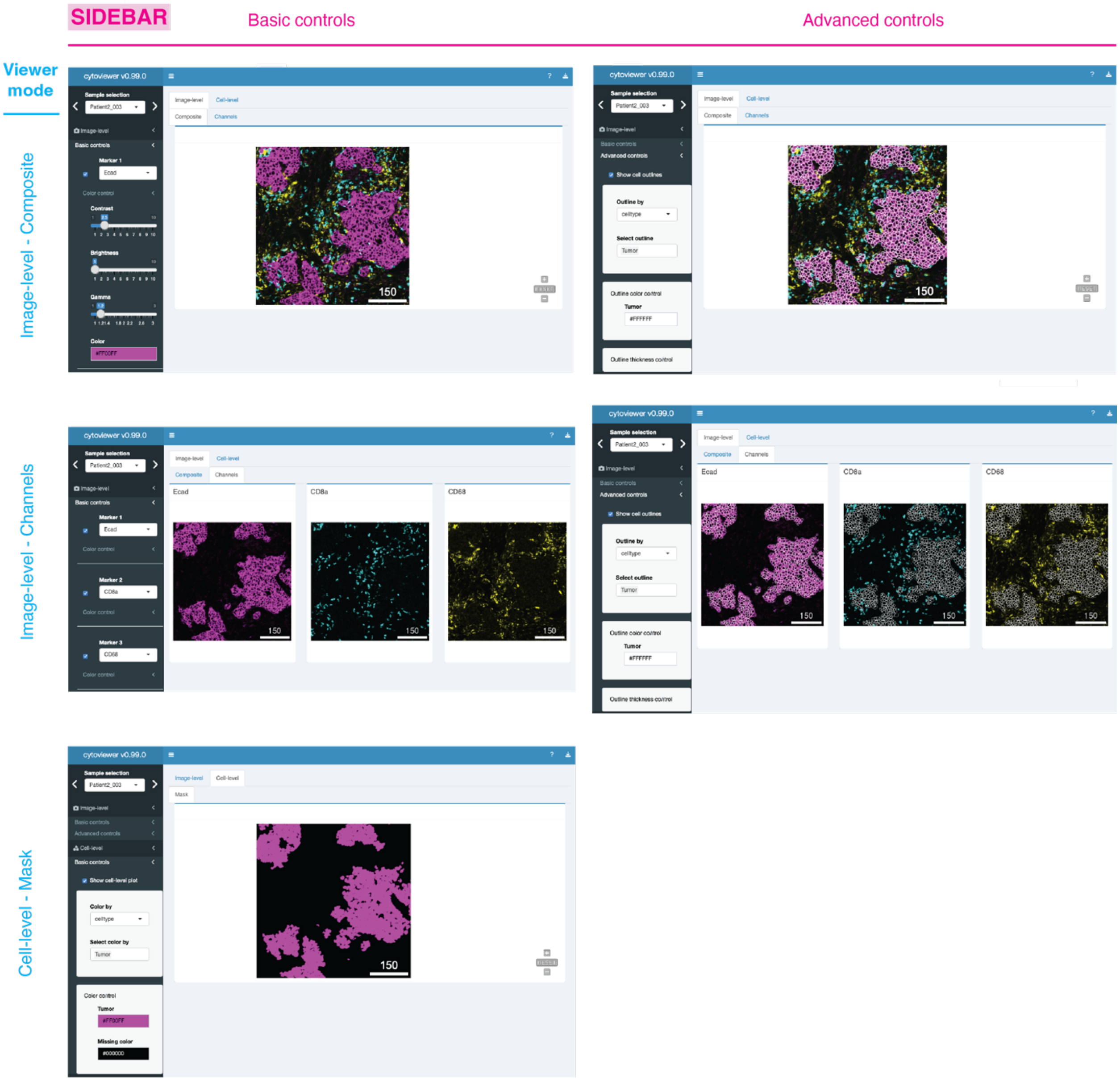


**Supplemental Figure S1: cytoviewer graphical user interface overview.**
The graphical user interface of cytoviewer for the three different viewer modes. Image-level-Composite with basic controls (top-left) and advanced controls (top-right), Image-level-Channels with basic controls (middle-left) and advanced controls (middle-right) and Cell-level-Mask with basic controls (bottom-left) are shown. For image-level visualization, Ecad (magenta), CD8a (cyan) and CD68 (yellow) marking tumor cells, CD8+ T cells and myeloid cells, respectively, are shown and channel color settings are as follows for all markers: Contrast: 2,5; Brightness: 1; Gamma: 1.2. For cell-level visualization, tumor cells (magenta) are highlighted. Note that the Composite and Mask tabs are zoomable. Scale bars: 150 $\mu$m.

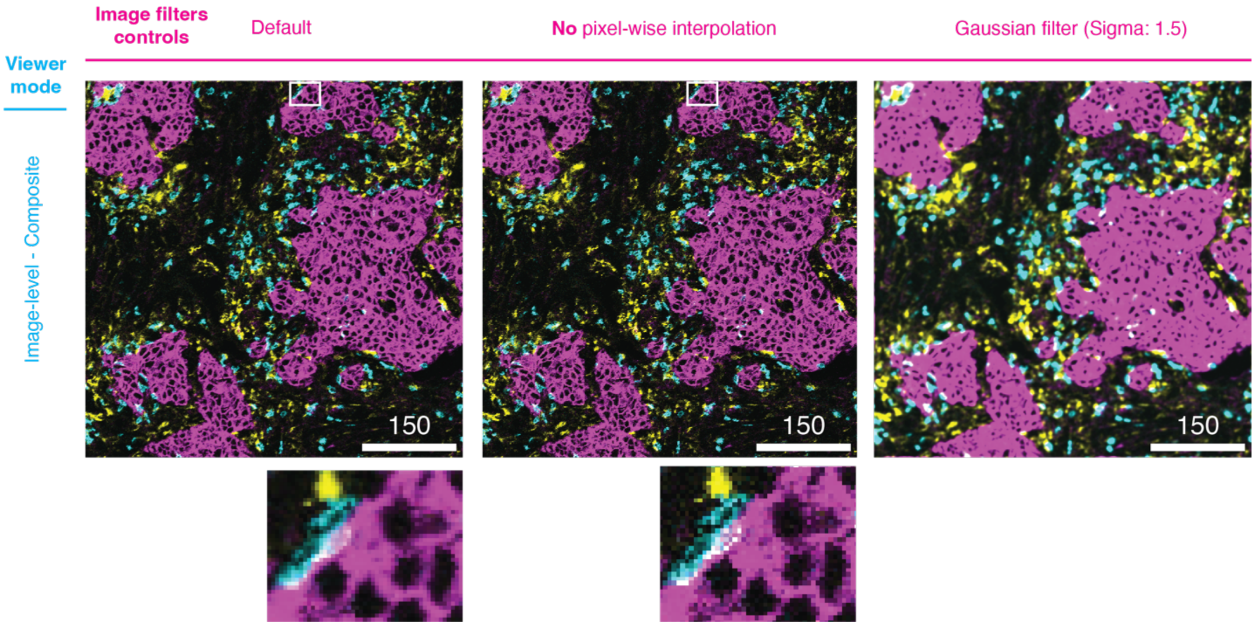


**Supplemental Figure S2: cytoviewer image filters.** Image filter controls are relevant for the image level (here: Composite). Ecad (magenta), CD8a (cyan) and CD68 (yellow) marking tumor cells, $CD8^+$ T cells, and myeloid cells, respectively, are shown. Channel color settings are as follows for all markers: Contrast: 2,5; Brightness: 1; Gamma: 1.2. The user can turn on pixel-wise interpolation (left, default) and off (center). The white boxes indicate the areas magnified in lower images. Users can also apply a Gaussian filter to the image (right, sigma: 1.5). Scale bars: 150 $\mu$m.

### S4 References


Eling, N., Damond, N., Hoch, T., & Bodenmiller, B. (2020). cytomapper: an R/Bioconductor package for visualization of highly multiplexed imaging data. *Bioinformatics*, *36*(24), 5706–5708. https://doi.org/10.1093/bioinformatics/btaa1061

Righelli, D., Weber, L. M., Crowell, H. L., Pardo, B., Collado-Torres, L., Ghazanfar, S., Lun, A. T. L., Hicks, S. C., & Risso, D. (2022). SpatialExperiment: infrastructure for spatially-resolved transcriptomics data in R using Bioconductor. *Bioinformatics*, *38*(11), 3128–3131. https://doi.org/10.1093/bioinformatics/btac299

Windhager, J., Bodenmiller, B., & Eling, N. (2021). An end-to-end workflow for multiplexed image processing and analysis. *BioRxiv*, 2021.11.12.468357. https://doi.org/10.1101/2021.11.12.468357
