## Supplementary material for "*cytoviewer:* an R/Bioconductor package for interactive visualization and exploration of highly multiplexed imaging data": R script: cytoviewer_publication_analysis.Rmd

````
---
title: "cytoviewer_publication_analysis"
author: "Lasse Meyer"
date: '2023-04-18'
output: html_document
editor_options: 
  markdown: 
    wrap: 72
---

```{r setup, include=FALSE}
knitr::opts_chunk$set(echo = TRUE)
```

# cytoviewer publication

R markdown file related to:

**cytoviewer: an R/Bioconductor package for interactive visualization
and exploration of highly multiplexed imaging data**

**Lasse Meyer, Nils Eling, Bernd Bodenmiller**

*Software note:* All analysis were performed using Bioconductor 3.16, R
version 4.2.1 (2022-06-23) and cytoviewer version 0.99.3.

# Analysis workflow

## Download example dataset

In the present manuscript we use an example imaging mass cytometry (IMC)
cancer dataset to showcase the functionality of `cytoviewer`. This
dataset was generated as part of the Integrated iMMUnoprofiling of large
adaptive CANcer patient cohort project ([immucan.eu](immucan.eu)) and
includes IMC data for 4 cancer patients diagnosed with different tumor
types (head and neck cancer, breast cancer, lung cancer and colorectal
cancer). Of note, the manuscript focuses on visualizing one breast
cancer image (Patient2_003).

The data input objects were processed with the [IMC data analysis
workflow](https://bodenmillergroup.github.io/IMCDataAnalysis/) and can
be downloaded from <https://zenodo.org/record/7079294>.

```{r download}
# Download - Note: Files are saved to the current working directory
download.file("https://zenodo.org/record/7647079/files/spe.rds",
              destfile = "spe.rds")
download.file("https://zenodo.org/record/7647079/files/images.rds",
              destfile = "images.rds")
download.file("https://zenodo.org/record/7647079/files/masks.rds",
              destfile = "masks.rds")
```

## Read data into R

### Images

The image data was stored as a `CytoImageList` object containing the
spillover corrected multi-channel images (n=14). Each image contains 40
channels and each channel represents the pixel-intensities of one marker
(proteins for IMC). The proteins for this dataset are immuno-oncology
related targets including Ecad, CD8a and CD68, which mark tumor, CD8+ T
cells and myeloid cells, respectively.

```{r images}
# Load images
images <- readRDS("images.rds")
images
```

### Segmentation masks

The segmentation masks were again stored as a `CytoImageList` object containing
one mask (n=14) for each image. Segmentation masks are defined as one-channel
images containing integer values for cells and zero for background.

```{r masks}
# Load masks
masks <- readRDS("masks.rds")
masks
```

### Metadata object

The metadata object was stored in `SpatialExperiment` format. It
contained various metadata information in the `colData` slot generated
during the analysis pipeline including patient-level information (such
as indication) and cell-level information (such as cell type and cell
area).

```{r spe}
# Load spe
spe <- readRDS("spe.rds")
spe
```

For more details regarding the input objects, please refer to the [IMC
data analysis
workflow](https://bodenmillergroup.github.io/IMCDataAnalysis/).

## Load cytoviewer

```{r load}
# Install cytoviewer 
if (!requireNamespace("BiocManager", quietly = TRUE))
    install.packages("BiocManager")

BiocManager::install("cytoviewer")

# Load cytoviewer
library(cytoviewer)
```

## Run cytoviewer

The `cytoviewer` function takes up to **five arguments** and the
functionality depends on which input objects are provided. For more
information, please refer to the help page found at `?cytoviewer` and
**Figure 1B** of this manuscript.

Here, we showcase the full functionality of `cytoviewer` by providing
the images, segmentation masks and metadata object that were described
above.

To match information between the `CytoImageList` objects (images, masks)
and the `SpatialExperiment` object (spe), two additional spots are
specified:

*img_id*: a single character indicating the `colData` (of the
`SpatialExperiment` object) and `elementMetadata` (of the
`CytoImageList` object) entry that contains the image identifiers. These
image ids have to match between the `SpatialExperiment` object and the
`CytoImageList` objects.

*cell_id*: a single character indicating the `colData` entry that
contains the cell identifiers. These should be integer values
corresponding to pixel-values in the segmentation masks.

```{r run}
# Create shiny app with cytoviewer
app <- cytoviewer(image = images, 
                  mask = masks, 
                  object = spe, 
                  img_id = "sample_id", 
                  cell_id = "ObjectNumber")

# Launch app in your web browser
if (interactive()) {
  shiny::runApp(app, launch.browser = TRUE)
  }
```

### Publication figure settings

To re-create the figures of the present manuscript, please follow the
instructions below **in order**:

#### Figure 1

**Figure 1A (Step 1)**:

Viewer mode: Image-level - Composite

Sample selection: Select *Patient2_003*

Image-level - Basic controls:

Select *Ecad* (magenta), *CD8a* (cyan) and *CD68* (yellow) with
following color control settings: Contrast: 2,5; Brightness: 1; Gamma:
1.2

**Figure 1C**:

**Top (Step 2)**:

Change Viewer mode: Image-level - Channels

**Middle/Left (Step 3)**:

Change Viewer mode: Image-level - Composite

Image-level - Advanced controls: Show cell outlines - Outline thickness
control: Select thick

**Middle/Center (Step 4)**:

Image-level - Advanced controls: Show cell outlines - Select *area* for
*Outline by*\` - Choose *plasma* for outline color control

**Middle/Right (Step 5)**:

Image-level - Advanced controls: Show cell outlines - Select *celltype*
for *Outline by* - Choose *white* for outline color control

**Bottom/Left (Step 6)**:

Change Viewer mode: Cell-level - Mask

Cell-level - Basic controls: Show cell-level plot

**Bottom/Center (Step 7)**:

Cell-level - Basic controls: Show cell-level plot - Select *area* for
*Color by* - Choose *plasma* for color control

**Bottom/Right (Step 8)**:

Cell-level - Basic controls: Show cell-level plot - Select *celltype*
for *Color by* - Choose *magenta* for tumor color control and *black*
for missing color

**Figure 1D**:

**Top (Step 9)**:

Go back to the settings at *Step 1*.

General - Basic controls - Image appearance: Change *Scale bar length*
to 100 and tick *Show Legend* and *Show Title*

**Bottom (Step 10)**:

Go back to the settings at *Step 8*.

Cell-level - Basic controls: Show cell-level plot - Select *celltype*
for *Color by* - Choose *Tumor*, *CD8* and *Myeloid* for *Select color
by* - Choose *magenta*, *cyan*, *yellow*, *black* for tumor, CD8,
myeloid and missing color color control

General - Basic controls - Image appearance: Change *Scale bar length*
to 100 and tick *Show Legend* and *Show Title*

#### Supplemental Figure 2

**Left (Step 11)**:

Go back to the settings at *Step 1*.

**Center (Step 12)**:

General - Basic controls - Image filters: Un-tick *Pixel-wise
interpolation*

**Right (Step 13)**:

General - Basic controls - Image filters: Tick *Gaussian filter* and set
it to 1.5

### Example cytomapper function usage

The R/Bioconductor `cytoviewer` package builds on top of the
R/Bioconductor `cytomapper` package
(<https://bioconductor.org/packages/release/bioc/html/cytomapper.html>)
and utilizes its functions and data containers.

Therefore, we can re-create images that were interactively generated
with `cytoviewer` using `cytomapper`. For more information refer to
**Supplementary Note S1.1**.

```{r load cytomapper}
# Install cytomapper
if (!requireNamespace("BiocManager", quietly = TRUE))
    install.packages("BiocManager")

BiocManager::install("cytomapper")

# Load cytomapper 
library(cytomapper)
```

Here, we showcase one example for **Figure 1A** of the manuscript using
the `plotPixels` function from `cytomapper`.

```{r run cytomapper}
# Re-create Figure 1A - Composite
plotPixels(image = images["Patient2_003"],
           colour_by = c("Ecad", "CD8a", "CD68"),
           colour = list(Ecad = c("black", "magenta"), 
                         CD8a = c("black","cyan"),
                         CD68 = c("black", "yellow")),
           bcg = list(Ecad = c(1, 2.5, 1.2), 
                      CD8a = c(1, 2.5, 1.2),
                      CD68 = c(1, 2.5, 1.2)),
           legend = NULL,
           image_title = NULL,
           scale_bar = list(length = 150)
           )   
```

```{r sessionInfo}
sessionInfo()
```
````
